# Metabolic evolution of pyranopterin-dependent biochemistry

**DOI:** 10.1101/2023.09.01.555371

**Authors:** Joshua E. Goldford, Ranjani Murali, Joan Selverstone Valentine, Woodward W. Fischer

**Affiliations:** Division of Geophysical and Planetary Sciences, California Institute of Technology, Pasadena, CA 91125, USA; Physics of Living Systems, Massachusetts Institute of Technology, Cambridge, MA 02139, USA; Blue Marble Space Institute of Science, Seattle, Washington, USA 98154; Department of Chemistry & Biochemistry, UCLA, Los Angeles, CA 90095, USA

**Keywords:** Ancient metabolism, biogeochemistry, evolution, metalloenzymes, protein language models, molybdenum, tungsten

## Abstract

Molybdenum (Mo)-dependent biochemistry is essential for many key metabolic pathways. However, theory and geological evidence suggests that its solubility during long intervals with low dioxygen would have limited its availability on early Earth. We developed models of metabolic evolution and found that reactions employing tungsten (W)-dependent biochemistry likely preceded Mo-dependent reactions, where Mo-usage increased dramatically after the production of dioxygen. Consistent with this finding, we analyzed genomes from over 65,000 phylogenetically diverse microbes and metagenomes from an environmental dataset, and we observed that dioxygen-utilizing prokaryotes living in aerobic niches are enriched with Mo-dependent enzymes as compared to anaerobic microbes. As an independent evaluation of this hypothesis, we combined protein language models, machine learning, and phylogenomic analysis to build a classifier for W- or Mo-pterin dependence in the DMSO reductase superfamily, and we found that W-pterin-dependent enzymes cluster near the root of the tree and that a subset of late-evolving aldehyde oxidoreductases (AORs) from aerobes are predicted to rely on Mo instead of W. Overall, our combination of metabolic modeling, phenotypic analysis, machine learning, and phylogenomic analysis suggest that Mo-pterin-dependent biochemistry likely derived from W-pterin-dependent biochemistry, and that Mo-usage increased drastically after the rise of oxygen.

## Introduction

Metals enhance cellular metabolism by providing functional diversity that far surpasses what organic macromolecules can achieve on their own; they play key catalytic roles in redox and acid-base reactions, the structuring of water, and in the transport of and manipulation of electrons and dioxygen. Consequently, it is estimated that over one quarter of all proteins are expected to bind a metal (1). The usage of metals in extant biology is driven by both mechanistic constraints as well as historical contingency, which is the outcome of dynamic interactions between time-varying environmental availability, evolving biochemical demands, and physiological flexibility (2).

The metal concentrations in surface environments have changed dramatically over Earth history as a function of the differential solubility, complexation, and redox behavior of distinct metals—modulated by vast state changes in the redox state of Earth’s surface environments and geochemical budgets set by different rock-forming minerals within the crust (2–5). For example, in the late Archean Eon, iron was abundant in seawater due to the lack of free dioxygen which allows ferrous iron to remain soluble (6). However, following the emergence of oxygenic photosynthesis, dioxygen produced by Cyanobacteria led to the oxidation of ferrous to ferric iron (7), causing it to precipitate out of seawater and form deposits of iron oxide, significantly reducing its availability in the oceans. In contrast, other metals like copper that are hosted in and form insoluble sulfide- and disulfide-phases, became solubilized after the oxygenation of the atmosphere, increasing global availability in seawater (2).

It is widely appreciated that the availability of molybdenum (Mo) would have been strongly influenced by the presence of dioxygen, where concentrations may have only reached ∼100 nM levels sometime after the Great Oxygenation Event (GOE) (8). This is in stark contrast with Mo-usage in biological systems today, where many diverse extant enzymes, including the most widespread (and potentially ancient) isozyme of nitrogenase (9–11), rely on a Mo-containing cofactor. While its widespread usage among many enzyme-catalyzed reactions has led to proposals that Mo was an essential component of the earliest forms of life (12, 13), other hypothesis suggest that similar metals (e.g. tungsten (W)) preceded Mo in the earliest phase of biochemical evolution (14). In modern biochemistry, Mo is found in either the complex FeMo cofactor (FeMoco) in Mo-dependent nitrogenase, or complexed to a dithiolene ligand from pyranopterin (typically called molybdopterin). In the former, FeMoco plays a unique and direct role in dinitrogen splitting and reduction, while in the latter the molybdopterin is responsible for facilitating a broad spectrum of metabolic reactions involving O-transfer (15). While FeMoco is unique to nitrogenases, molybdopterin is used by a wide number of reactions and is a critical component in the active sites of diverse enzymes like DMSO reductases, sulfite oxidase, formate dehydrogenase, xanthine oxidase and nitrate reductase, among others (14). For a subset of pyranopterin-dependent enzymes, W is either required or substitutable in lieu of Mo, where it facilitates the same O-transfer chemistry (16). For example, several aldehyde reductases such as aldehyde:ferredoxin oxidoreductase (AOR) and glyceraldehyde:ferredoxin oxidoreductase (GAPOR) depend on tungstopterin instead of molybdopterin cofactors, and are often found in hyperthermophilic archaea (16). Modern day usage of Mo may give the perception of a critical dependence throughout the history of life, but *in vitro* enzymology studies indicate that Mo and W can be substituted in many enzymes (14, 16–19), raising the possibility that the dependency of specific metals in ancient pyranopterin-dependent enzymes could have been significantly different than what appears today in modern enzymes. Taken together, while Mo is widely used in extant biological systems, it remains unclear whether this pattern reflects primordial use, or alternatively if Mo was preceded by W or a similar metal during evolutionary history.

Physicochemical differences between W and Mo likely led to distinct responses to environmental changes throughout Earth’s history. W and Mo have differential environmental availability despite having similar abundance in Earth’s crust (20): Mo is often much more abundant than W in oxygenated marine systems (21), but these differences decrease in sulfidic and anoxic (euxinic) conditions (22). In hydrothermal systems, W can even exceed Mo concentrations (23) . These differences in environmental abundances are likely due to several factors, including the higher reduction potential of molybdenum relative to tungsten compounds or increased likelihood of thiomolybdate formation and precipitation (24–26). The VI to IV reduction potential is typically higher for Mo than for W regardless of the ligand system (27), indicating that in ecosystems with low environmental reduction potential (*E*_h_), Mo(VI) may be reduced to Mo(IV), while W(VI) remains oxidized. Equilibrium models of tungstate and molybdate/molybdenum sulfides suggests the selective depletion of molybdate at concentrations of sulfide between ∼10 uM to 1 mM, while tungsten disulfides will form only in environments with >1 mM sulfide in marine environments (24–26), suggesting that tungstate may have been more available in both the ferruginous Archean and plausible euxinic Phanerozoic marine environments than in modern day oceans. Additionally, the lower reduction potentials of W complexes compared to Mo suggest that W-pterin may be the preferred cofactor for biochemical reactions involving lower potentials, which have been hypothesized to be more common in the earliest phases of metabolic evolution (14, 16).

In this paper, we explored the hypothesis that tungstopterin preceded molybdopterin in metabolic evolution. We examined the biological record using a suite of independent approaches, leveraging comparative genomics, environmental metagenomics in a redox-stratified environment, metabolic network modeling, machine learning and phylogenomics to characterize molybdopterin and tungstopterin cofactor usage throughout the biosphere and evaluate evolutionary trajectories using a recently developed model of metabolic evolution. We found that tungstopterin-dependent reactions are predicted to have emerged before the emergence of oxidative photosynthesis, while the majority molybdopterin-dependent reactions emerged after oxygenic photosynthesis. We also found a strong association between metal preference in pterin-dependent genes and O_2_ physiology and aerobic environmental niches across phylogenetically diverse microbes and that tungstopterin-dependent genes appear preferentially near the root of the DMSO reductase superfamily. Finally we used machine learning to predict metal cofactor preferences for over 20,000 orthologs of aldehyde:ferredoxin oxidoreductase (AOR) and find a subset of orthologs from aerobic prokaryotes that are predicted to bind Mo. We demonstrate that key evolutionary questions can be addressed by combining metabolic network evolutionary modeling, phylogenomics and machine learning, enabling self-consistent modeling for biochemical evolution through Earth history.

## Results

### A multidisciplinary approach to inference of metabolic evolution and ecology

To explore the evolution of pyranopterin biochemistry, we combine several approaches, quantifying molybdopterin and tungstopterin usage in modern biology and exploring evolutionary models of metabolic and protein evolution. First, we address the question of whether molybdopterin and tungstopterin usage is driven by environmental redox gradients and physiological factors in modern day microbes and ecosystems using comparative biology techniques and environmental metagenomics. Second, we leverage evolutionary models of metabolic network evolution, machine learning and phylogenomic analysis to ordinate molybdopterin and tungstopterin-usage throughout evolutionary history. To first access the relationship between environmental availability of Mo and W, and the biological demand for Mo and W, we first explore how pyranopterin-dependent genes change in a well characterized environment with a strong redox gradient and variable Mo/W availability.

### Quantifying the abundance of tungsto- and molybdopterin in the Black Sea

We sought to determine to what degree environmental availability of Mo/W in natural environments shaped potential biological utilization by analyzing metagenomes from the water column of the Black Sea. Geochemical modeling suggests that Mo and W should have strong differential availabilities as a function of dioxygen and sulfide concentrations in the environment (Fig 1a, Methods). Under sufficiently sulfidic environments, thermodynamic modeling suggests that oxygen fugacity predominantly controls the speciation between soluble oxyanions (e.g. MoO_4_^2-^) versus insoluble metal disulfides (e.g. MoS_2_). The Black Sea is a meromictic basin with a steep chemocline with a sharp drop in dioxygen around ∼100 m in depth, and an increasing rise in sulfide concentrations throughout the water column (Fig 1b). Previous studies have measured dissolved and particulate Mo/W in depths up to 2000 m (22) and have found a negative relationship between depth and Mo concentration and a positive relationship in dissolved W concentration and depth (Fig 1d).

**Fig. 1:**
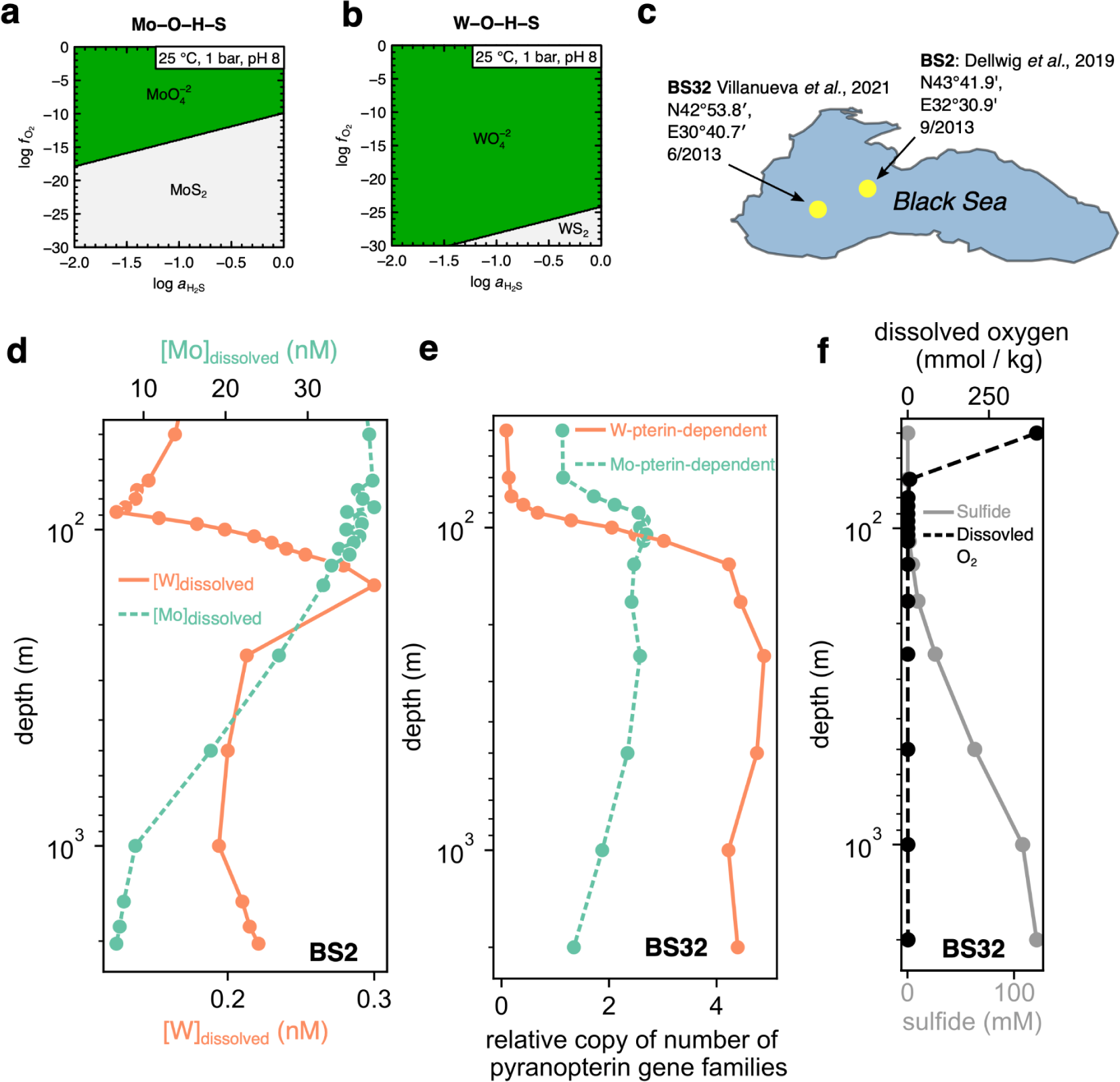
Pyranopterin-utilizing gene abundance in the Black Sea is associated with metal availability. **(a)** Phase plot showing the energetically dominant species for Molybdenum (Mo) oxyanions and sulfides under different hydrogen sulfide activities (*x*-axis) and dioxygen fugacities (*y*-axis). Green regions indicate the soluble oxyanion phase while gray regions denote the disulfide (MoS_2_) phase. (**b**) same as **a**, but for Tungsten (W). **(c)** sampling locations for dissolved oxyanions (BS2) and metagenomics samples (BS32) in the western gyre of the Black Sea. **(d)** Dissolved tungstate (orange line) and molybdate (green line) in the black sea water column (from sampling station BS2, (22)). **(e)** mean copy numbers of either Mo-dependent (green line) or W-dependent (orange line) pyranopterin-binding gene families (Supplementary Table S1) from 15 metagenomic samples in the Black sea western gyre (NCBI BioProject PRJNA649215) sampling site BS32. **(f)** dissolved dioxygen (black line) and sulfide (grey line) in the from sampling station BS32.

To examine whether changes in the availability of these trace metals were reflected in the ecosystem-level biochemistry, we analyzed shotgun metagenomics from microbial communities sampled at 15 different depths along the water column (28). We annotated 160 metagenome-assembled genomes (MAGs) assembled from these data with all pyranopterin-binding gene families in the Kyoto Encyclopedia of Genes and Genomes (KEGG) (*n*=41, Supplementary Table S1-2). Relative abundances of these MAGs were quantified, allowing us to compute the relative copy number of genes from orthogroups that use either molybdopterin or tungstopterin (Supplementary Table 3, see Methods). We found that the average copy number of W-pterin-dependent genes per genome sharply increased below the oxycline and maintained a high copy throughout the deep sulfidic water column (Fig. 1e-f). In contrast, Mo-pterin-dependent genes are prevalent in surface waters, peak just above the oxycline and slightly decrease with depth in the water column (Fig. 1e-f). To probe further this trend in apparent metal usage, we computed the copy numbers of the tungstate-specific transporters TupABC per genome and found that *tupA*, *tupB* and *tupC* all increased with depth throughout the water column—in line with increase in tungstopterin-dependent genes (Supplementary Figure S1).

These results suggest that dioxygen availability may be positively associated with ecosystem-level Mo-pterin-dependent biochemistry, and negatively associated with W-pterin-dependent biochemistry. To determine whether or not this relationship could more generally shape genomic structure on evolutionary time-scales (rather than just gene functional structure on ecological time-scales), we sought to explore the possibility that microbial genomes with known O_2_ requirements may more generally encode preferences for either Mo or W-pterin-dependent biochemistry.

### Aerobes are enriched with Mo-pterin biochemistry

Our analysis of the Black Sea metagenomes suggest that dioxygen availability may control ecosystem-level Mo-pterin-dependent biochemistry differently than W-pterin-dependent biochemistry. We hypothesized that dioxygen requirements of prokaryotes may be predictive of Mo or W-dependent biochemistry, where aerobes would be more dependent on Mo-dependent enzymes than anaerobes, and anaerobes would be more dependent on W-dependent enzymes than aerobes. To explore this possibility, we computed the proportion of genes that were Mo-pterin and W-pterin-dependent per genome across a diverse group of prokaryotes spanning the tree of life. We identified all KO groups for 65,703 representative species from the Genome Taxonomy Database (29) (GTDB, version r207) using the KEGG specific HMM profiles and cutoffs (see Methods), and computed the proportion of KO groups per genome that bound to either Mo-pterin or W-pterin (*n*=41, Supplementary Table S1). For each species cluster in the GTDB, we obtained dioxygen sensitivity data (30), and analyzed all taxa labeled as either obligate aerobe (anaerobe) or aerobe (anaerobe) (see Methods) (Supplementary Table S4). Consistent with our expectations, we observed that aerobes are heavily enriched with Mo-pterin-dependent gene families relative to anaerobes (Fig. 2a, Mann-Whitney *U* test: *P* < 10^-78^). We additionally found that anaerobes are enriched with tungstopterin-dependent enzymes compared to aerobes (Fig. 2b, Mann-Whitney *U* test: *P* < 10^-63^).

**Fig. 2:**
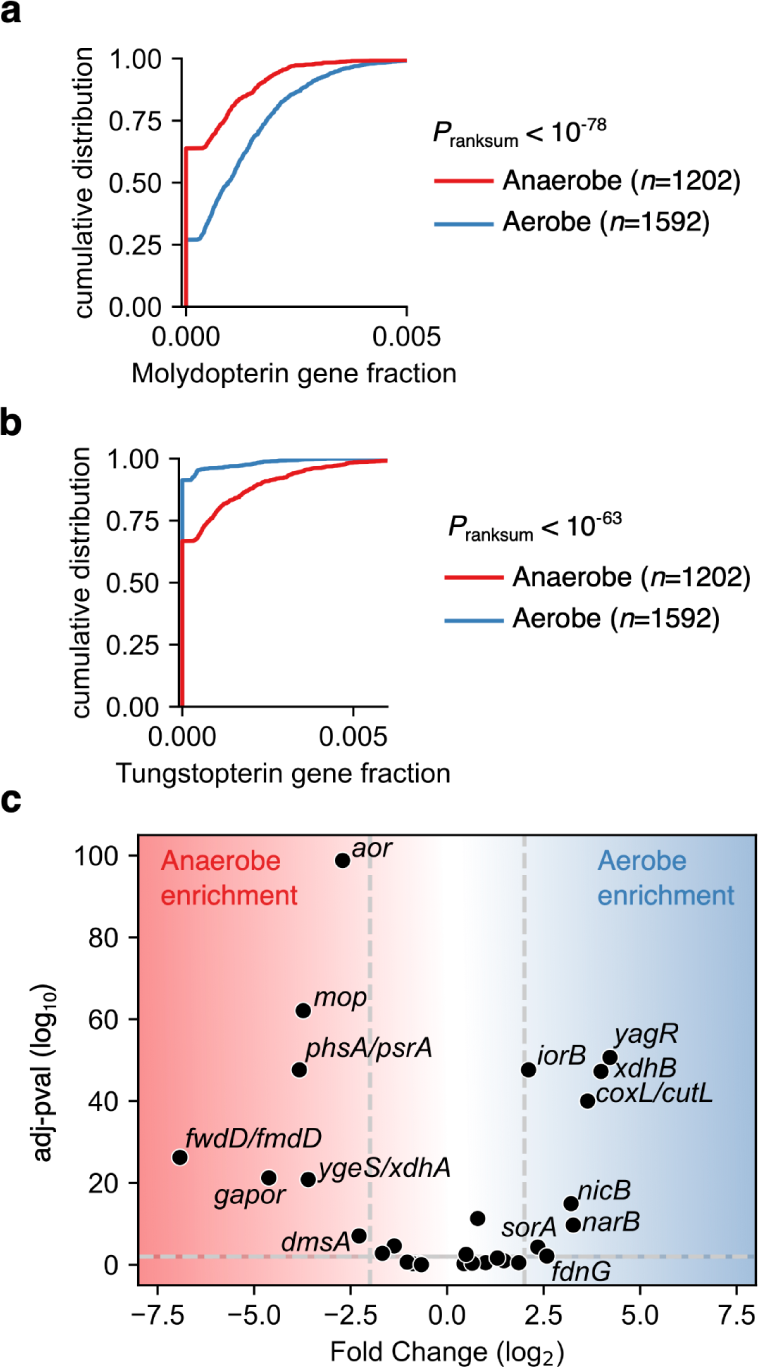
Aerobic prokaryotes are enriched with molybdopterin whereas anaerobes are enriched with tungstopterin-utilizing genes. **(a-b)** We annotated 65,703 representative species from the Genome Taxonomy Database (GTDB) (29) with KEGG (KO) orthogroups, and computed the proportion of genes in each genome mapped to a molybdopterin (or tungstopterin) binding domain (Supplementary Table S1). We used datasets that had classified prokaryotes as either aerobes or anaerobes using the JGI GOLD metadata (30), and plotted the empirical cumulative distribution (*y*-axis) plots of molybdopterin (tungstopterin) gene fraction (*x*-axis). (**c**) a volcano plot showing the fold change of gene family copy number between aerobes and anaerobes (*x*-axis) plotted vs. the adjusted *p*-value (y-axis) after multiple comparisons correction (see Methods).

To determine which gene families were enriched in aerobes or anaerobes, we generated a volcano plot (Methods); for each pterin-binding gene family we plotted the average fold change in relative gene copy number in aerobes vs. anaerobes against a measure of statistical significance (Fig. 2c). We found that tungstopterin-dependent enzymes like aldehyde:ferredoxin (*aor*), glyceraldehyde-phosphate:ferredoxin (*gapor*) were all highly enriched in anaerobes.

Formylmethanofuran dehydrogenase (*fwdD/fmdD*) were also highly enriched in anaerobes, as well as thiosulfate/polysulfide reductase (e.g. *phsA*/*psrA*). Consistent with expectations, aerobes are much more enriched with the aerobic carbon-monoxide dehydrogenase large subunit (*coxL*/*cutL*). We also found that the pyranopterin binding domains for ferredoxin-nitrate reductase (*narB*), nicotinate dehydrogenase subunit A (*nicA*) and sulfite dehydrogenase (cytochrome) (*sorA*) were all enriched in aerobes compared to anaerobes. Interestingly, anaerobes and aerobes have different preferences for gene families that encode the pyranopterin-binding submit for similar reactions: Aerobes are enriched with the *yagR* xanthine dehydrogenase (K11177), while anaerobes tend to have higher copy numbers of the *ygeS*/*xdhA* (K00087).

The strong relationship between dioxygen availability and the pyranopterin metal dependence within microbial communities in the Black Sea and the association between oxygen requirement and W vs Mo-pterin usage in phylogenetically diverse microbes suggest that dioxygen may drive the differences in availability and utilizing preference between Mo and W-pterin across both ecological and evolutionary time-scales. Since O_2_ availability is one of the few atmospheric variables to change throughout deep time (31), we explored the hypothesis that W-pterin biochemistry emerged prior to Mo-pterin biochemistry using a recently developed model of planetary-scale metabolic evolution (32).

### W-pterin-dependent reactions emerge earlier than Mo-pterin dependent biochemistry

To analyze the usage of pyranopterin cofactor dependence throughout metabolic evolution, we traced the usage of molybdopterin (tungstopterin) coenzyme-dependent reactions using metabolic network expansion (33–36). The network expansion algorithm simulates the emergence of metabolic networks by determining which biochemical reactions are possible given a set of starting, “seed” molecules. These reactions then produce products, which are added to the seed compound set, and new reactions are now feasible with the additional set of compounds. This process is repeated until no more additional compounds can be added to the network. This computational technique has been used to address a variety of questions, including how dioxygen affected the biosphere (34) and whether phosphate was essential in the earliest metabolic networks (36, 37) . Recently, network expansion was used to uncover a plausible evolutionary trajectory of core metabolism (32). This trajectory was generated by performing network expansion on an manually curated planetary-scale network of metabolic reactions, considering both thermodynamic and catalytic constraints. The starting compound set includes metals and inorganic metal species (e.g. molybdate, tungstate), as well as reduced nitrogen (ammonia), sulfur (sulfide), orthophosphate, and 21 organic molecules produced in simple prebiotic chemistry experiments (38). The inclusion of a non-ATP dependent purine biosynthesis pathway was necessary to achieve expansion from this seed set, and it produced a trajectory of metabolic network evolution where the production of quinones resulted in the emergence of oxygenic photosynthesis in the later stages of the expansion (32) (Fig 3b-c).

**Fig. 3:**
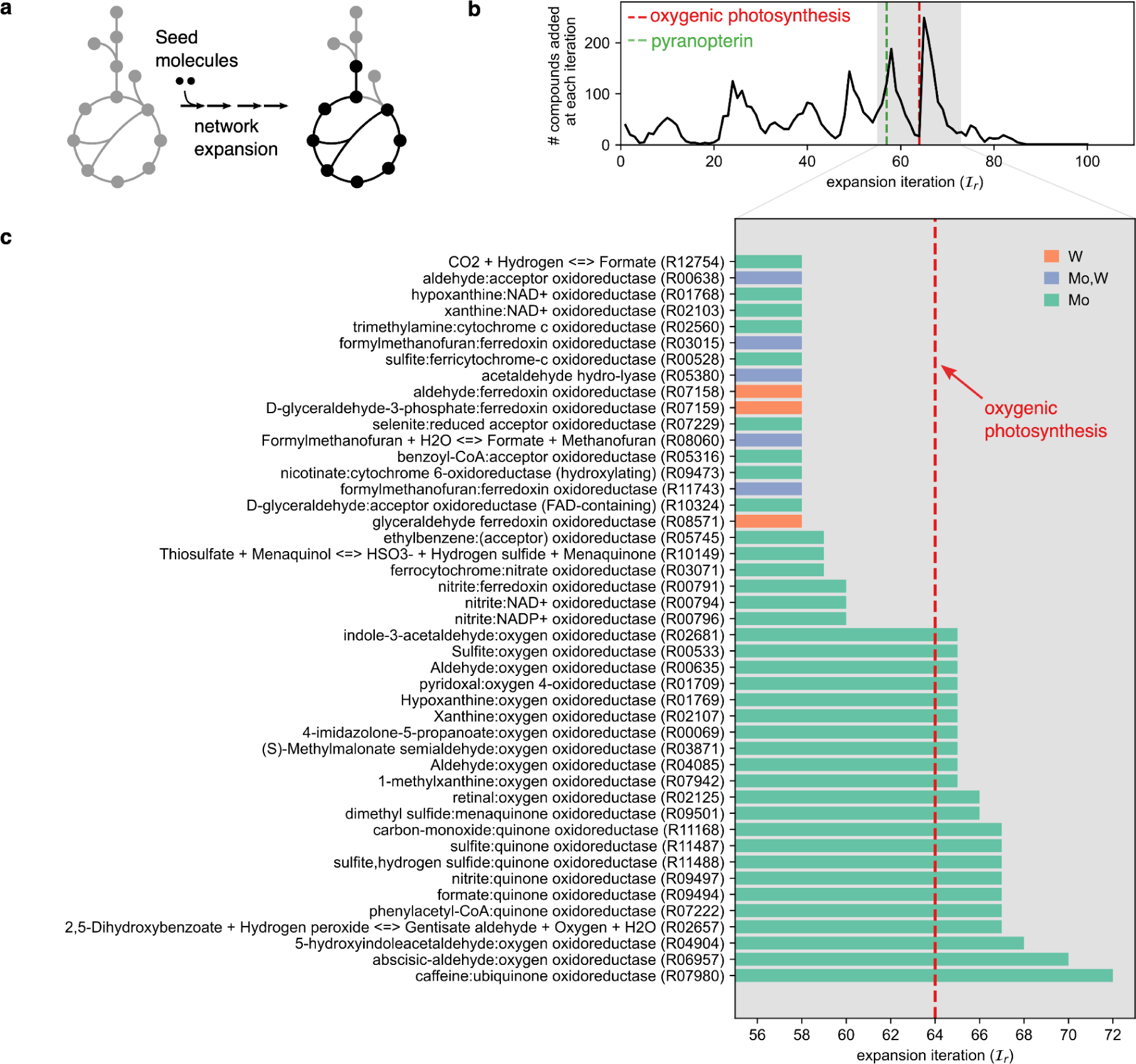
Tungstopterin-utilizing reactions tend to emerge before Molybdopterin-dependent reactions throughout the model of network evolution. **(a)** We analyzed a recent model of biochemical evolution using network expansion (32). **(b)** We performed network expansion as done previously, and plotted the number of compounds produced at each expansion iteration (*x*-axis). The horizontal green line denotes when pyranopterin emerged during network expansion. The gray shaded region denotes the expansion region where pyranopterin-dependent reactions are recruiting into the expanding network. **(c)** a horizontal bar plot showing iteration of emergence for each pyranopterin-dependent reaction in our model (*n*=45). The color of the bar denotes the particular metal dependence of the reaction, where reactions can either depend on Mo (green), W (orange), or either Mo or W (purple). All reactions that can use W emerge before oxygenic photosynthesis (red dotted line).

We first analyzed the expansion iterations where either molybdopterin-dependent reactions or tungstopterin-dependent reactions emerged throughout the expansion. We found that ∼41% (15/37) of the molybdopterin-dependent reactions emerge before oxygenic photosynthesis, whereas 100% (8/8) of the tungstopterin-dependent (or reactions than can use tungstopterin) reactions emerged before dioxygen (Fig 3c, orange and purple bars). Note that several of the molybdopterin-dependent reactions that emerged after oxygenic photosynthesis did not use O_2_ directly as a substrate (*n*=10). Rather, these reactions operated on reaction products that only emerged after O_2_ (e.g. xanthoxin dehydrogenase (E.C. 1.1.1.288)) or relied on quinones (e.g. phenylacetyl-CoA:quinone oxidoreductase (see Supplementary Table S5).

Molybdopterin-dependent reactions that emerged before oxygenic photosynthesis were several cytochrome-dependent or ferredoxin-dependent oxidoreductases operating on oxidized sulfur or nitrogen species (e.g. nitrate reductase and sulfite dehydrogenase) (see Supplementary Table S5). Interestingly, all the molybdopterin-dependent reactions using iron-dependent coenzymes (ferricytochrome and ferredoxin) emerged prior to oxygenic photosynthesis, while most quinone-dependent transformations occurred after oxygenic photosynthesis, except for the menaquinone-dependent thiosulfate reductase (E.C. 1.8.5.5, see Supplementary Table S5).

To examine whether the difference between the emergence time for W-pterin dependent reactions versus Mo-pterin-dependent reaction was primarily driven by the differential usage before and after dioxygen is generated within the expansion, we performed 10^4^ simulations where we randomly sampled 10 compounds as additional seed molecules with and without O_2_, and performed network expansion (see Methods). We found that the mean expansion iteration for W-pterin-dependent reactions was smaller than for Mo-pterin-dependent reactions when O_2_ was generated during the expansion. However, when O_2_ was added as a seed molecule, the mean expansion iterations for W and Mo-pterin-dependent reactions were comparable (Supplementary Fig. S2).

All together, results from metabolic expansion suggest that W-pterin biochemistry emerged, on average, earlier than Mo-pterin-dependent reactions. We next sought to explore whether signals of this evolutionary trajectory for pterin-dependent biochemistry may be present in the phylogenomic record.

### Phylogenomics support for early W-pterin usage in DMSO reductase family

Our results from metabolic modeling, metagenomic analysis and analysis of pyranopterin-dependent biochemistry among prokaryotes have suggested that tungstopterin-dependent biochemistry preceded molybdopterin biochemistry, potentially before the oxygenation of atmosphere via oxygenic photosynthesis. We sought to examine whether these results could be corroborated by molecular phylogenomics techniques. Recent work has suggested that newly developed machine learning techniques from natural language processing (39) can be leveraged to predict non-trivial structural and functional features from protein sequences (40, 41). Recent success of such models prompted us to build a classifier for pyranopterin-binding domains by first embedding sequences using a transformer model and training a regularized logistic regression classifier on sequences annotated as binding either tungstopterin or molybdopterin (Fig. 4a, see Methods). Stratified-cross validation based on sequence similarity revealed that the trained classifier had ∼99% accuracy (Methods, Supplementary Figure S3). We applied this classifier to all 132,260 protein sequences in the GTDB mapping pyranopterin-binding gene families (Methods) and computed the proportions of each gene family that were predicted to either bind W or Mo-pterin (Fig. 4b). This model predicted the majority of anaerobic aldehyde ferredoxin oxidoreductases (GAPOR and AOR) sequences were tungstopterin-specific, while the majority of all other gene families were predicted to use Mo-pterin. We next applied our classifier to a set of sequences from the DMSO reductase family (*n*=1568), which had previously been used to construct a maximum likelihood phylogenetic tree (42) (Fig. 4c, Supplementary Table S6). Consistent with our expectation, the root of the tree consisted primarily of sequences from *aor* and *gapor* gene families predicted to use tungstopterin. However, a minority of predicted tungstopterin-binding variants of major gene families like *phsA*/*psrA* and *narG* were predicted to be derived from Molybdopterin-specific gene families. These results suggest that while the majority of tungstopterin sequences are predicted to be ancestral for the DMSO reductase family, several tungstopterin-specific enzymes may be evolutionarily derived in a minority of instances.

**Fig. 4:**
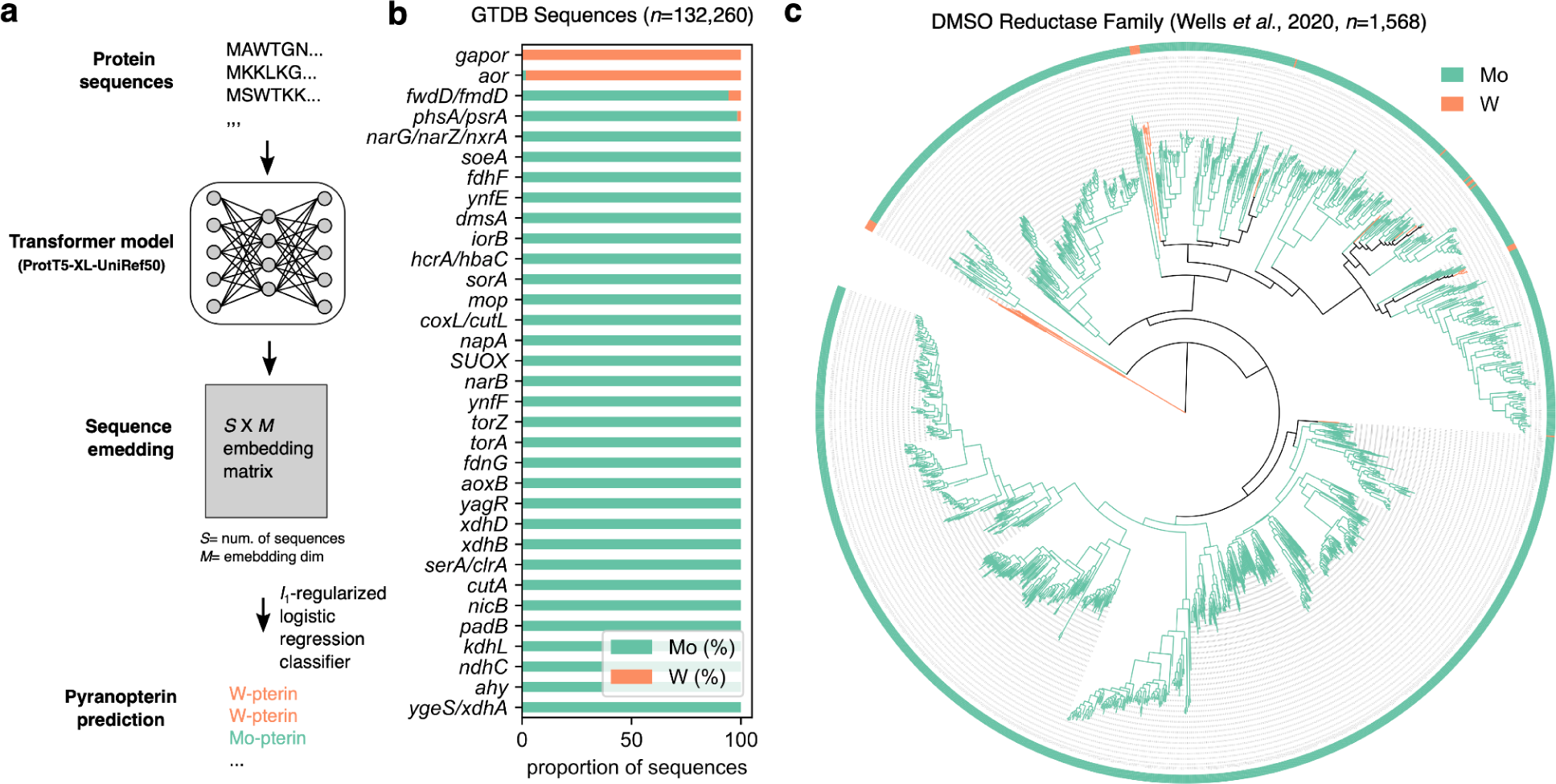
Predicting molybdopterin or tungstopterin specificity of enzymes using protein language models. **(a)** We built a machine learning classifier to predict tungstopterin or molybdopterin. We downloaded *n*=1003 sequences from the Uniprot database annotated to bind either tungstopterin (CHEBI: 60215) or molybdopterin (CHEBI: 25372) (see Methods), and used the ProtT5-XL-Uniref50 protein language model (40) to create 1024-dimensional vector embeddings for each protein sequence. We trained a logistic regression model classifier with *l*_1_ regularization and cross-validation to predict the cofactor specificity using the embedding vector, revealing ∼99% cross validation accuracy (see Supplemental Figure S3). **(b)** Predicted pyranopterin preferences for pterin-binding domains from 41 gene families (KEGG orthologous groups, see Supplementary Table S1) from the GTDB. The proportion of each gene family (*y*-axis) that was predicted to bind either Mo-pterin (green) or W-pterin (orange) are plotted as a stacked bar plot. **(c)** Maximum likelihood tree of the DMSO reductase family (42), colored by the predicted metal preference.

### Machine learning reveal derived Mo-dependent orthologs from the aldehyde:ferredoxin oxidoreductases

We used our machine learning classifier to predict metal preference for all orthologs mapping to the aldehyde:ferredoxin oxidoreductase gene family (K03738) from representative genomes in the GTDB (*n*=23,308). Surprisingly, we found that 1.67% (*n*=388) were predicted to use Mo, while the rest were predicted to be dependent on W (Supplementary Table S7). We hypothesized that these putative Mo-dependent AORs may have derived from the more common W-dependent variants throughout evolutionary history as response to oxygenation of the atmosphere. To explore this possibility, we took a subset of these sequences (*n*=1195) that derived from genomes with known physiological dependencies for O_2_ (see Methods), and constructed a maximum likelihood phylogenetic tree (see Methods). Phylogenetic analysis suggests that all Mo-dependent sequences are likely derived from W-dependent ancestors (Fig 5a). Consistent with our hypothesis, we found that Mo-dependent AORs are almost exclusively found in aerobes: 44/48 of the Mo-dependent AORs are derived from aerobes (Fisher’s exact test: *P* < 10^-26^) (Fig 5b, Supplementary Table S7).

**Fig. 5:**
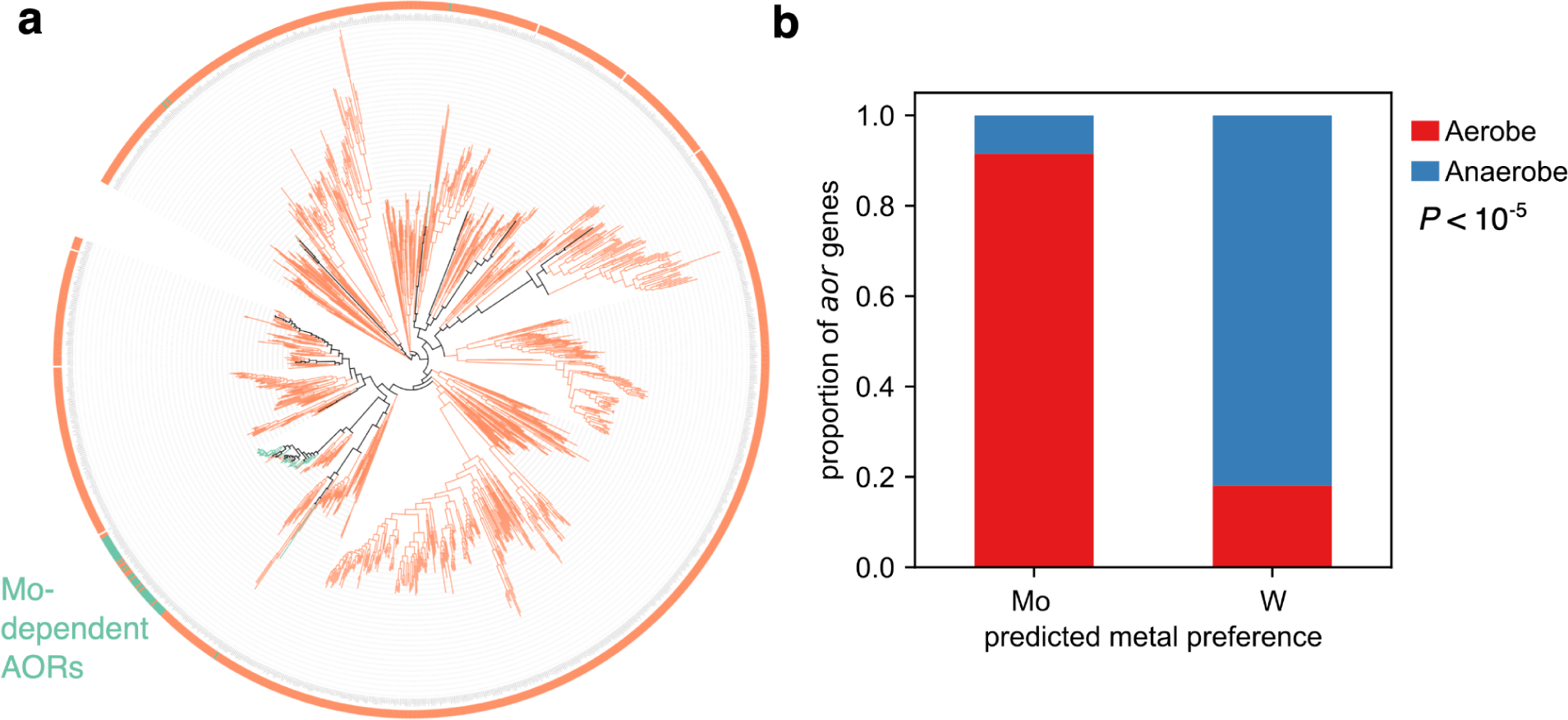
Protein language models predict an evolutionarily-derived Mo-dependent aldehyde:ferredoxin oxidoreductase. **(a)** A maximum likelihood tree for orthologs mapping to the aldehyde:ferredoxin oxidoreductase gene family (K03738) from representative genomes in the GTDB with oxygen physiology annotations (*n*=1,195). The tree was rooted with minimal ancestor deviation (MAD) rooting (52), and tips with light green (orange) represent predicted Mo-pterin dependent (W-pterin dependent) AORs. **(b)** A stacked bar chart composed of the proportion of predicted Mo or W-pterin dependent AORs from aerobes (red) and anaerobes (blue).

Mechanistic studies explore the possible energetic difference between Mo and W in formaldehyde:ferredoxin oxidoreductase in *Pyrococcus furiosus* have shown that the metalation with Mo (instead of W) renders the enzyme inactive (43). Quantum chemical studies have indicated that this is due to the fact that formation of the terminal oxo intermediate is endergonic because of the low reduction potential of the ferredoxin electron acceptor (44). Thus, if these predicted Mo-dependent AORs are active, they would likely require a stronger oxidant than the low potential ferredoxins observed in the canonical W-dependent AORs.

## Discussion

In this study, we investigated the hypothesis that tungstopterin preceded molybdopterin in metabolic evolution. Our results provide multiple lines of independent evidence supporting this hypothesis. By employing a comprehensive suite of methods exploring both the extant biosphere as well as plausible evolutionary trajectories, our findings indicate that molybdopterin-dependent reactions predominantly emerged following the rise of oxygenic photosynthesis, in contrast to tungstopterin-dependent biochemistry. The association between metal preference in pterin-dependent enzymes and O_2_ physiology across diverse microbes further supports this temporal differentiation in cofactor utilization. By harnessing the power of metabolic network evolutionary modeling, phylogenomics, and machine learning, we demonstrated that key questions in the field of biochemical evolution can be addressed using multiple lines of evidence, paving the way for future studies aimed at uncovering the intricate history of metabolic evolution through Earth’s history.

While the history of metal usage in the biosphere has traditionally been examined through the lens of geochemical availability (45, 46) or metabolic demand (46), our results suggest that biochemical constraints, specifically in relation to available redox coenzymes and cofactors, might have influenced the evolution of metal usage. For Mo and W in the context of pyranopterin-dependent reactions, our model indicates that pyranopterin is formed relatively late in metabolic evolution (Fig 3). Additionally, our findings indicate that even if molybdate was more abundant in Archean or Phanerozoic oceans, the preference for Mo might have only materialized after both the emergence of pyranopterin and high-potential electron acceptors, such as quinones, became prevalent in the biosphere. Our data from genome composition (Fig 2a-b), metabolic network evolution (Fig 3a-b), and environmental metagenomics (Fig 1) emphasize that dioxygen availability seems to be the dominant environmental variable shaping the metal preference in pyranopterin-containing enzymes. Notably, the vast majority of pyranopterin-dependent reactions utilizing high-potential acceptors, like quinones and O_2_, exclusively employ Mo (Fig 3, Supplementary Table S5). Conversely, reactions involving oxidants with lower redox potential, such as oxidized ferredoxins, exclusively utilize W (Fig 3, Supplementary Table S5). This observation aligns with prior associations between redox potentials and metal specificity (16, 17, 47), and underscores the intricate relationship between the redox potential of involved biochemical partners and the evolution of metal cofactor usage in these enzymes.

The hypothesis that Mo usage was preceded by W throughout the history of the biosphere not only has implications for the evolution of pyranopterin-dependent enzymes, also may suggest that ancestors of the modern molybdenum-iron nitrogenases were originally dependent on W, and that tungsten-iron nitrogenase may even exist in the extant biosphere. Phylogenomic analysis and ancestral sequence reconstruction suggests that extant vanadium-iron and iron-iron nitrogenases derived from sequences similar to modern molybdenum-iron nitrogenases before the GOE (9–11). While it is tempting to hypothesize that these ancestral nitrogenases may have relied on W instead of Mo, no evidence to date suggests that molybdenum-iron nitrogenases can function when substituted with W. For instance, biophysical studies of a W-substituted nitrogenase from the phototrophic non-sulfur bacterium *Rhodobacter capsulatus* suggest that FeWco redox potential was significantly lower than FeMoco, which prevents reduction of FeWco during the reaction cycle (48). Thus, mechanistic constraints may have simultaneously favored both the usage of Mo in nitrogenase over W, and W over Mo in pyranopterin-dependent reactions early in biochemical evolution. That being said, experimental confirmation of the cofactor dependence in natural nitrogenases is highly underexplored, especially from anaerobes with distinct clades of nitrogenase enzymes (9).

Our work highlights the power of leveraging multiple analysis to uncover potential novel enzymes in the biosphere. In particular, we used machine learning models trained on metal preference for pyranopterin-dependent enzymes to uncover a small subset of aldehyde:oxidoreductases from aerobes that may be Mo-dependent. While a representative structure from AlphaFold2 (49) reveal a novel acidic loop near the proximal iron sulfur cluster and a conserved histidine near the active site (Supplemental Figure S4), it is unclear to what extent these features modify the reduction potential in such a way to lower the energy barriers of transition state. Future structural and quantum chemical modeling studies (44) of these predicted Mo-dependent AORs may quantitatively reveal the effect these active site differences may have on energetics during the reaction cycle.

Future work can expand on this work by exploring evolutionary history across many more sequences for Mo/W-dependent enzyme families (50), or by investigating other metals, such as copper or iron, with well known geological histories (45). However, much care must be taken into consideration as metals in enzymes can have many roles, some of which are substitutable generically (e.g. divalent cations) or under specific conditions (51). Overall, we expect that combining the power of comparative biology, phylogenomics, machine learning and metabolic network analysis has the potential to resolve the deepest puzzles in the history of the biosphere.

## Acknowledgements

J.E.G. acknowledges support by the Gordon and Betty Moore Foundation as Physics of Living Systems Fellows through grant number GBMF4513, Caltech Center for Comparative Planetary Evolution, and the Simons Foundation.

## Contributions

J.E.G., J.S.V., and W.W.F. designed the research. J.E.G. prepared data. J.E.G. wrote code and ran simulations. J.E.G. and R.M. performed analysis. J.E.G. wrote the manuscript. All authors read and approved the final manuscript.

## Materials and Methods

### Software availability

Code and data are available on the following github repository: https://github.com/jgoldford/pyranopterin

### Geochemical modeling

Phase plots for Mo and W species were generated using the CHNOSZ package in R (53) and SUPCRT92 (54). The phase diagrams were created under specific conditions set at a temperature of 25°C, a pressure of 1bar, and a pH of 8. The range for H_2_S activity was between 10^-2^ and 1, while the O_2_ activity ranged from 10^-30^ to 1. The activity for the species in the aqueous medium was set at 10^-8^. For the generation of these phase plots, the chemical basis was established using MoO_4_ ^-2^, H S, H O, dioxygen, and H^+^ for Mo species. Similarly, for W species, the basis was established using WO ^-2^, H S, H O, dioxygen, and H^+^. Subsequent to defining the chemical basis, species of interest were retrieved using specific criteria: species containing Mo or W along with O, H, and S that are either in aqueous or crystalline form. Using this information, affinity calculations were performed followed by the construction of the phase diagrams. Importantly, custom thermodynamic data for specific chemical species (WS_2_, WO_2_, and WO_3_) was obtained from the Geological Survey Bulletin 1259 (55).

### Genome annotation and physiology

Metagenome assembled genomes from the Genome taxonomy database (GTDB) v207 (29) or from the Black Sea (28) were annotated with Kofamscan (56) using default parameters. Only annotations to Kegg Orthologs (KO) that exceeded the default thresholds in Kofamscan were considered positive hits. Oxygen requirement annotations were obtained from a study which aggregated annotations from the Joint Genome Institute (JGI) Genomes Online Database (GOLD) (30). Taxa labeled as “obligate aerobe” or just “aerobe” were classified as aerobic, while taxa labeled “obligate anaerobe” or “anaerobe” were classified as anaerobe. NCBI taxonomy IDs were mapped to GTDB IDs using metadata provided on the FTP server from the GTDB. We identified KEGG orthologous group (KO groups) by first identifying KEGG reactions previously annotated as using tungstopterin or molybdopterin, and identifying the orthologous group of the pyranopterin subunit manually (see Supplementary Table S1).

### Metagenomic sequencing analysis

To determine the relative abundances of Metagenome-Assembled Genomes (MAGs) in metagenomic samples, a computational pipeline was devised using MAGs previously downloaded from the Sequence Read Archive (SRA) database. Each metagenomic sample underwent alignment to the MAGs reference using the bbmap.sh script. Post alignment, the outputs, in Sequence Alignment/Map (SAM) format, were filtered to retain only reads with a mapping quality of 10 or above. These filtered SAM files were subsequently converted to the binary BAM format using samtools view. The resultant BAM files were then sorted, and indices were generated for each one using samtools. Following this, indexed and sorted BAM files were used to produce statistical summaries with samtools *idxstats*, which was used to estimate the number of reads that aligned to each reference MAG. The relative abundance of each MAG in the metagenomic samples estimated by computing the fraction of reads that mapped to that MAG in each sample. The entire procedure was automated with a Python script, using tools such as samtools v1.8 and bbmap.sh, and performed on the Reznick high-performance computing cluster at Caltech.

To compute relative abundances of Mo or W-dependent genes from metagenomic sequences, we took the *n*=39 (5) gene families that could use Mo (W) in the pyranopterin, computed the total number of genes per MAG mapping to these gene groups. For each sample *k*, the average copy number of Mo-pterin and W-pterin genes per sample (*z’*_Mo, k,_ *z’*_W, k_) were computed as follows:

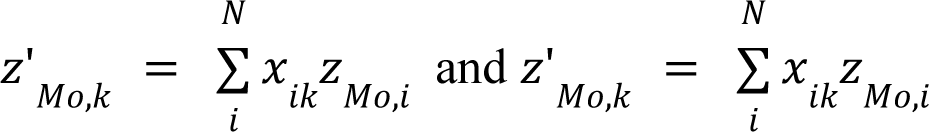

Where *x_ik_* is the relative abundance of MAG *i* in sample *k*, and *z_Mo,i_* and *z_W,i_* are the copy numbers of genes that use Mo-pterin and W-pterin, respectively, in MAG *i*.

### Network expansion

The network expansion algorithm was run as described previously (36, 37). Briefly, seed compounds are allowed to react given the reactions in the network, which then produce product compounds. The product compounds are added to the seed set, and the process is repeated until convergence. The network expansion algorithm was implemented using the `networkExpansionPy` python package (https://github.com/jgoldford/networkExpansionPy). The expansion trajectory was produced using the same seed compounds used previously. For 10^4^ expansions with randomly chosen seed sets (Supplementary Figure 2), we sampled 10 additional organic compounds and added them to the seed set. For each sample, we created a seed set both with and without dioxygen.

### Machine learning classifier for Mo/W dependence

We downloaded *n*=1003sequences from the Uniprot database annotated to bind either tungstopterin (CHEBI: 60215) or molybdopterin (CHEBI: 25372). For the Mo-dependent proteins, we only considered protein sequences from the SwissProt database (*n*=642), while for W-dependent proteins, we considered data from TrEMBL (*n*=361), due to the small number of manually reviewed tunstopterin-dependent enzymes in SwissProt (*n*=12). All proteins were downloaded directly from the Uniprot website.

We used a transformer-based T5 model (Rostlab/prot_t5_xl_half_uniref50-enc) from the Hugging Face model hub to generate protein sequences embeddings. Sequences were first processed to remove white spaces, gaps, and convert non-standard amino acids to characterize consistent with the tokenizer vocabulary. To compute embeddings, each protein sequence was tokenized and then passed through the T5 model to generate token-level embeddings. These embeddings are averaged across the entire protein sequence, producing a singular 1024-dimensional vector that represents the protein.

Using the sci-kit learn library, sequence clusters were employed for cross-validation analysis on cofactor predictions. We used CD-HIT (57) to cluster sequences at 80% sequence similarity, and partitioned *n=*1003 sequences into 416 clusters. For each sequence cluster, the sequences in the cluster constitute the test set, and the remaining sequences formed the training set. A logistic regression classifier from sci-kit learn, equipped with *L*_1_ regularization, was trained on this training data using the LogisticRegressionCV class. Features for the classifier were the 1024-dimensional embedding vector for each protein. The performance of this classifier was determined using balanced accuracy scores for both the training and test datasets.

### Phylogenetic analysis

1094 sequences of enzymes within the DMSO reductase superfamily in GTDB were combined with a reference sequence of aldehyde ferredoxin oxidoreductase from *Pyrococcus furiosus* (PDB id: 1AOR_1) and aligned using MUSCLE (58). This multiple sequence alignment was used to infer a phylogenetic tree using IQTREE-2 (59) with a Dayhoff substitution substitution matrix and 1000 ultrafast bootstraps. The phylogenetic tree was visualized using the Interactive Tree Of Life (iTOL) server (60).

### Statistical analysis

We performed statistical analyses using Python 3.6 with version 1.5.4 of the `scipy.stats` module. For comparing two groups in Fig. 2, we utilized the two-sided, nonparametric Mann Whitney *U* test, using the `mannwhitneyu` subroutine. To account for multiple comparisons, we applied the Benjamini-Hochberg correction for adjusting the p-values in the `statsmodels` library. Lastly, we performed a Fisher’s exact test using the `fisher_exact` function.

## Supplementary Materials

Supplementary Tables S1-7

Supplementary Figures S1-4

## Supplementary Figures

**Fig. S1:**
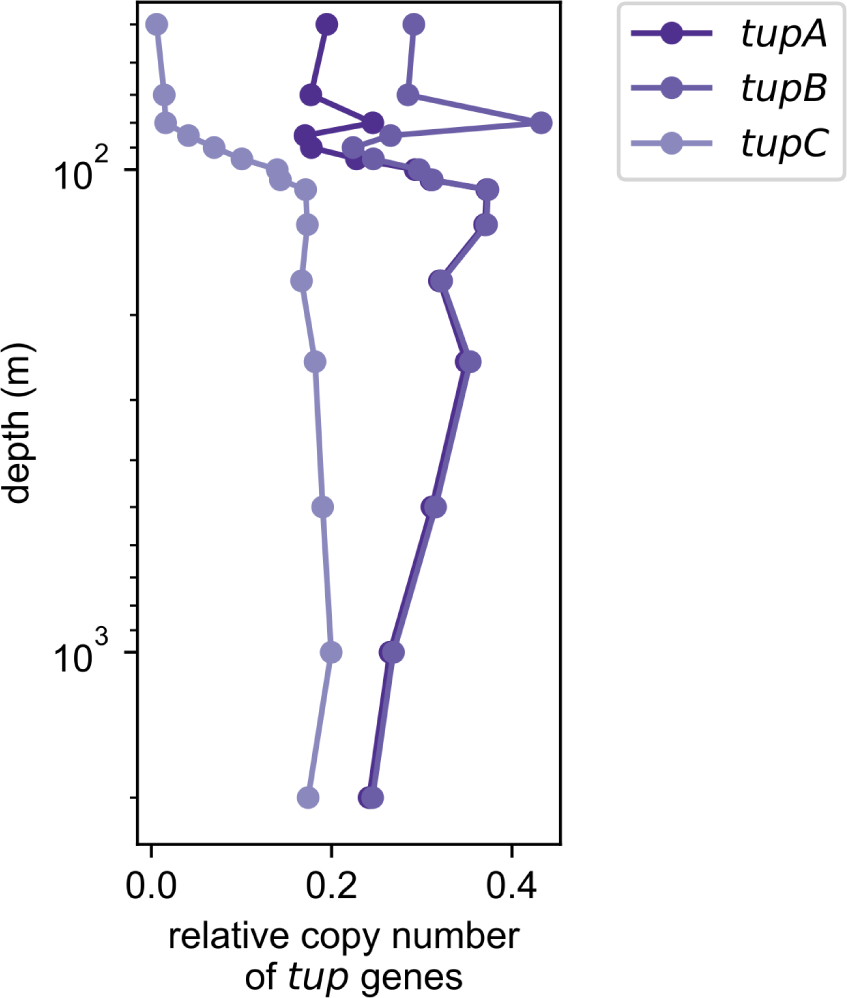
Relative copy numbers of tungstate transporters throughout the Black Sea. The relative copy numbers of *tupA* (K05772), *tupB* (K05773) and *tupC* (K06857) genes (dark, medium and light purple, respectively).

**Fig. S2:**
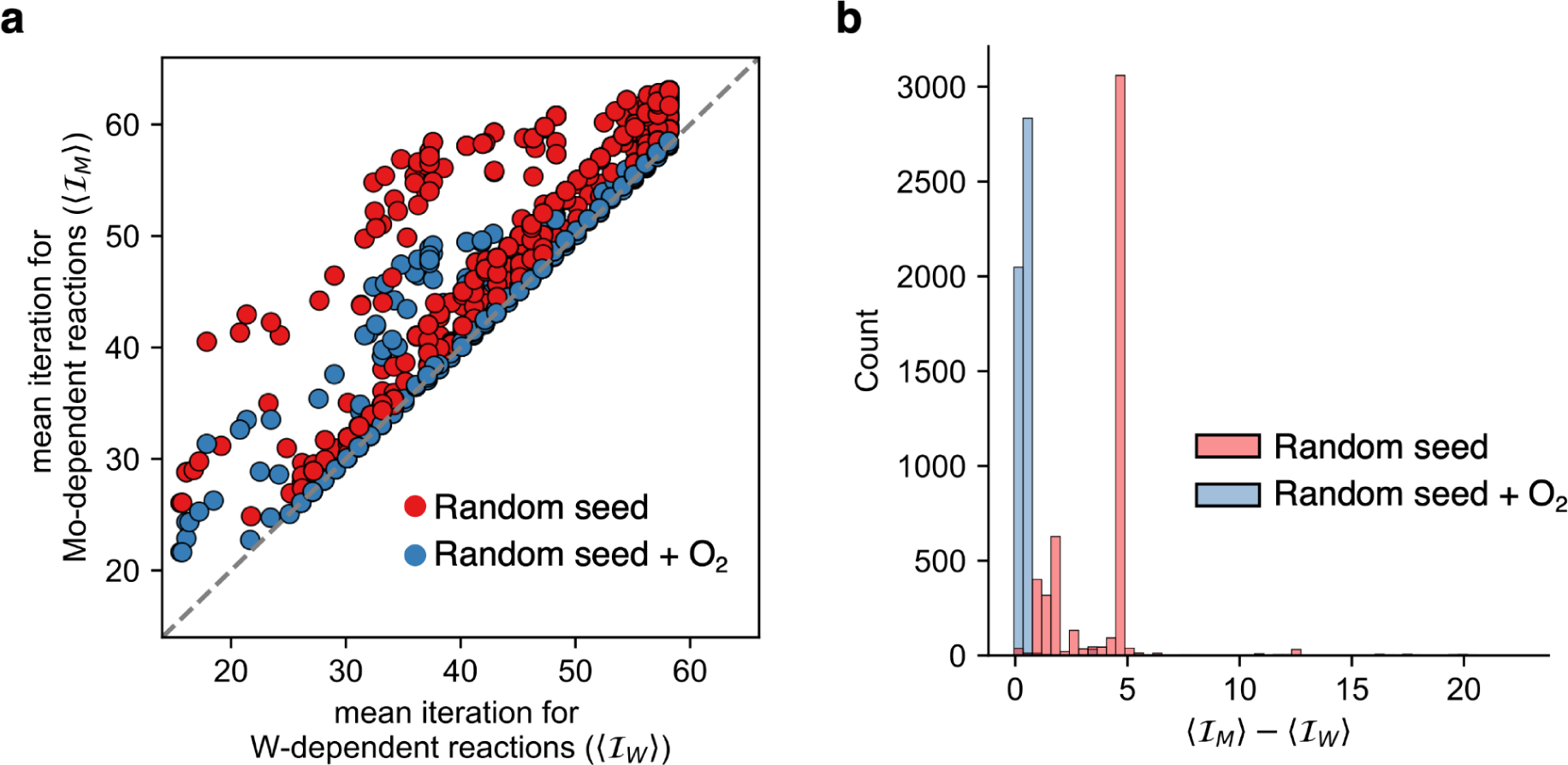
The emergence of Mo and W-dependent biochemistry is dependent on O_2_. **(a)** We performed 10^4^ simulations where we randomly sampled 10 compounds as additional seed molecules with and without O_2_, and performed network expansion (see Methods). For all Mo (W)-pterin dependent reactions, we computed the mean expansion iteration < 𝐼_𝑀𝑜_ > (< 𝐼_𝑊_ >) and plotted this value on the *y* (*x*) axis, where dots colored red were generated from seed sets without dioxygen, while blue indicates seed sets generated with dioxygen. In all cases, the mean expansion iteration for Mo exceeded or equaled W. **(b)** The difference between the mean expansion iteration between Mo-pterin dependent reactions and the W-pterin dependent reactions, colored by whether dioxygen was provided in the seed set. The difference between the expansions is strongly influenced by the presence of dioxygen, and is much larger when dioxygen is produced endogenously (via oxygenic photosynthesis) compared to when provided as a seed molecule.

**Fig. S3:**
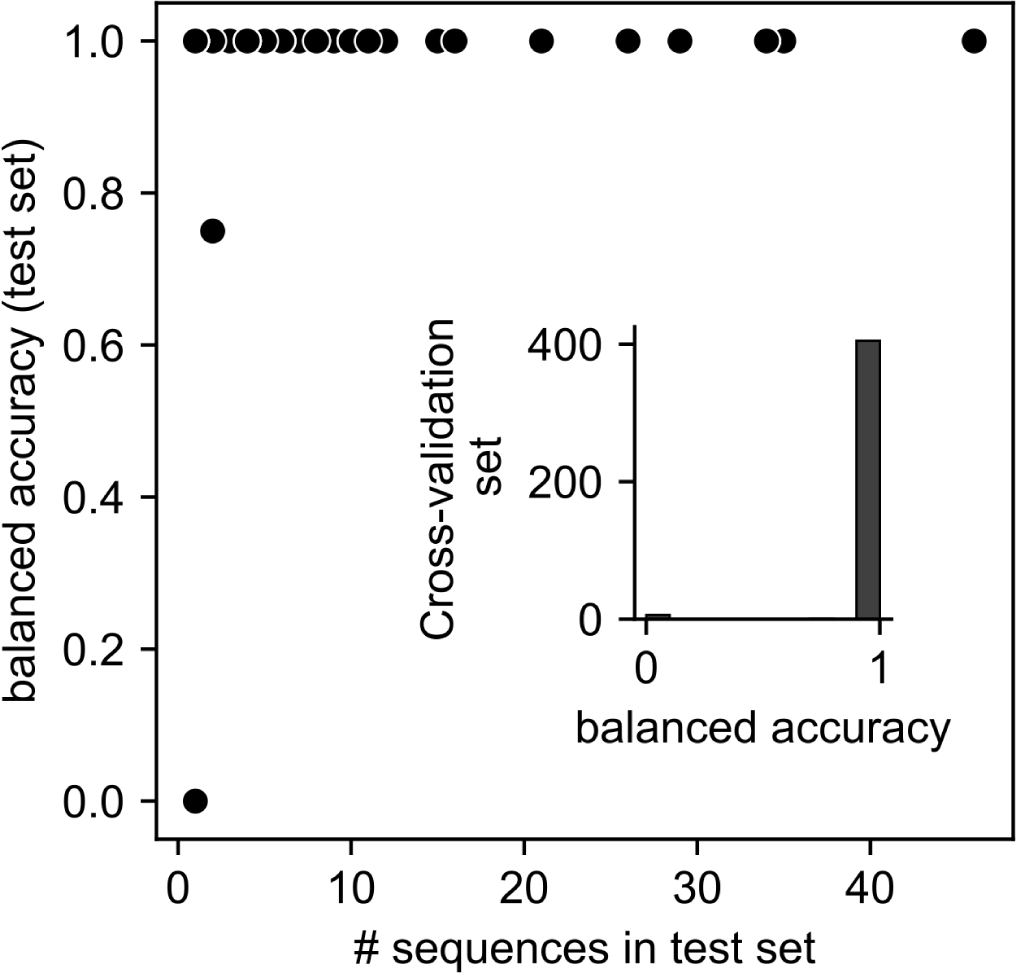
Cross-fold validation for pyranopterin metal dependence classifier. A scatterplot showing balanced accuracies in test sets (*y*-axis) vs. the number of sequences in the test set (*x*-axis). Cross fold validation was performed by clustering sequences at 80% sequence similarity, then holding out an entire cluster as a test set. (inset) A histogram of balanced accuracies for all 416 test sets.

**Fig. S4:**
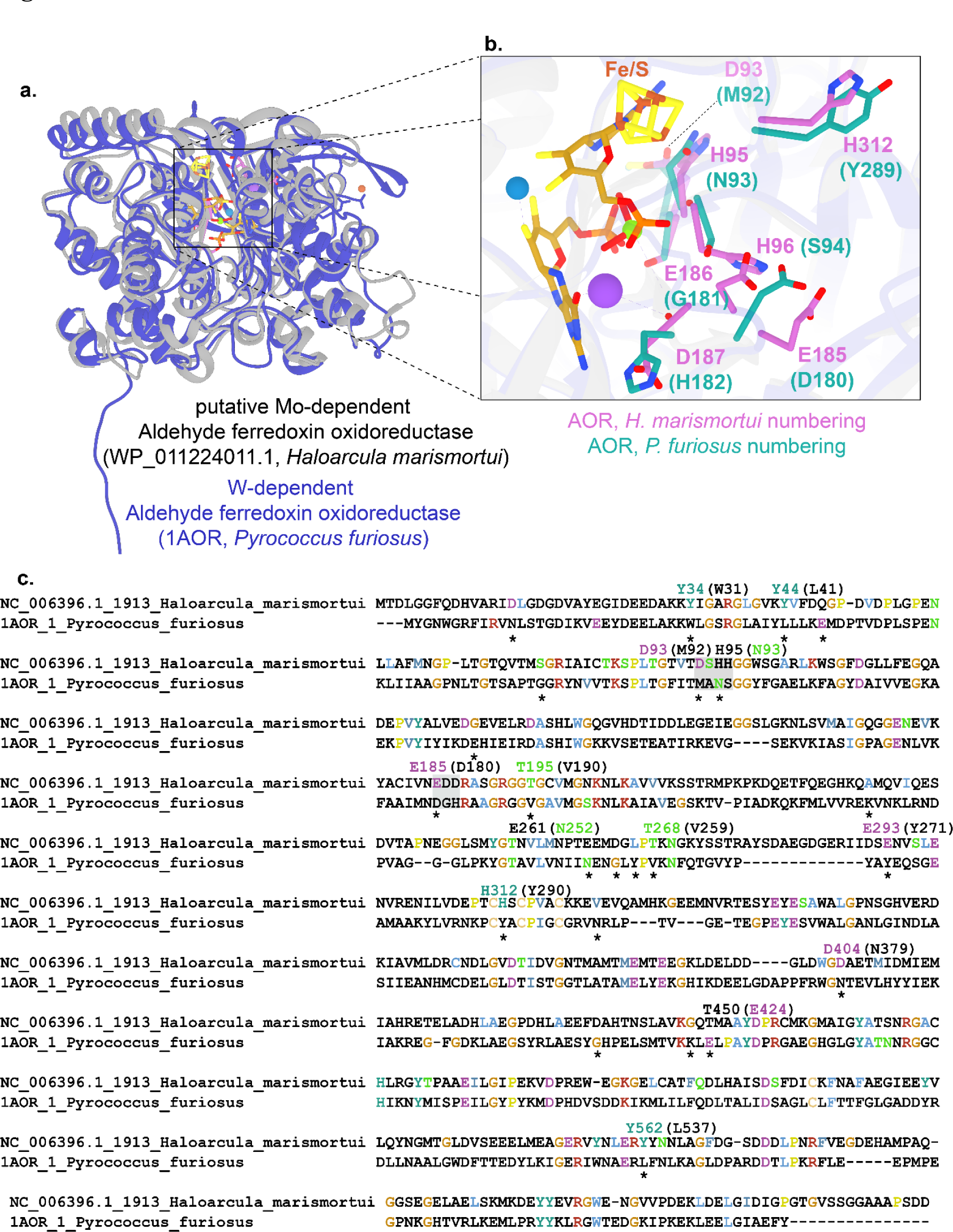
Structural model of putative Mo-dependent aldehyde oxidoreductase. **a.** Structural model generated by Alphafold2 of the aldehyde oxidoreductase from *Haloarcula marismortui* was download from the UniProt database (61) and superimposed with the crystal structure of the W-dependent AOR from *Pyrococcus furiosus* (PDB ID: 1AOR) with the use of the software, Chimera (62). **b.** Highly conserved residues (>98% conserved) adjacent to the Molybdopterin and Fe/S cluster containing the active site were colored. The highly conserved residues were conserved across the clades containing the putative Mo-dependent and W-dependent AORs, and were identified from the multiple sequence alignment depicted in c. **c.** Multiple sequence alignment of the aldehyde oxidoreductases generated by MUSCLE (58) with highly conserved residues (>98% conserved) colored using the CLUSTAL-based scoring matrix and identified from the multiple sequence alignment of the tree depicted in Figure 5, visualized using Jalview (63).

